# Inferring cell diversity in single cell data using consortium-scale epigenetic data as a biological anchor for cell identity

**DOI:** 10.1101/2022.10.12.512003

**Authors:** Yuliangzi Sun, Woo Jun Shim, Sophie Shen, Enakshi Sinniah, Duy Pham, Zezhuo Su, Dalia Mizikovsky, Melanie D. White, Joshua W.K. Ho, Quan Nguyen, Mikael Bodén, Nathan J. Palpant

## Abstract

Methods for cell clustering and gene expression from single-cell RNA sequencing (scRNA-seq) data are essential for biological interpretation of cell processes. Here we present TRIAGE-Cluster which uses genome-wide epigenetic data from diverse bio-samples to identify genes demarcating cell diversity in scRNA-seq data. TRIAGE-Cluster integrates patterns of repressive chromatin deposited across diverse cell types with weighted density estimation to determine cell type clusters in a 2D UMAP space. We then present TRIAGE-ParseR, a machine learning method that evaluates gene expression rank lists to define gene groups governing the identity and function of cell types. We demonstrate the utility of this two-step approach using atlases of *in vivo* and *in vitro* cell diversification and organogenesis. We also provide a web accessible dashboard for analysis and download of data and software. Collectively, genome-wide epigenetic repression provides a versatile strategy to define cell diversity and study gene regulation of scRNA-seq data.

## Introduction

Single-cell RNA sequencing provides unprecedented opportunities to study the diversity of cell processes underpinning development and disease. Standard analysis workflows involve stepwise analytic tools that decompose high dimensional data to reveal the underlying complexity of and relationships between cell types. However, the computational challenges involved in single cell analysis are significant at least in part because of common user-defined parameter settings that make interpretation of cell diversity a subjective exercise. Downstream analyses are also dependent on prior knowledge, relying on known marker genes, subjective data subtraction, and the use of reference annotation databases. Currently, few methods use unsupervised orthogonal reference points for defining cellular diversity and interpreting genetic regulation of cell states.

Cell type clustering methods underpin downstream analyses to guide hypotheses about cellular processes. Most current clustering algorithms group cells based on gene expression similarity, relying on mathematically defined constraints for data analysis [1]. Agglomerative clustering methods such as SINCERA [2], pcaReduce [3], and CIDR [4] use distance measurements such as Euclidean, Manhattan, Jaccard, Minkowski, and Canberra [5] to merge clusters iteratively based on similarities [6]. K-means clustering groups data points by taking a distance measure and iteratively optimising the centroid of each group to minimize within-cluster distance [7]. Model-based approaches like Gaussian Mixture Models (GMM) exemplified by scGMAI [8] use (multiple) distributions to capture relationships between cells [7]. Density-based clustering such as DBSCAN [9] and GiniClust [10] are non-parametric approaches that group cells based on data point density, where low-density areas are considered as outliers. Lastly, graph-based clustering is an extension of density-based clustering [11] where relationships among cells are represented by a similarity graph [12]. Louvain and Leiden are popular clustering approaches implemented in Seurat [13] and SCANPY [14].

Owing to the complex nature of cell clustering there is no generally accepted standard for parameter setting or method selection [1]. Furthermore, most methods assume all cells in a data set are meaningful and are therefore interpreted equally while it is well appreciated that sequencing depth, ambient RNA, and technical variation in scRNA-seq data are major variables influencing data variability and accuracy [15-17]. Outside of routine data trimming for doublets and high-quality cells, establishing justifiable and generalisable criteria for data exclusion remains challenging. Similarly, gene expression analysis often depends on arbitrary selection of computational parameters and reference points. For instance, differential expression gene analysis (DE), the use of highly variable genes [18] between cell clusters results in data interpretation relevant to individual data sets [19].

To address these limitations, newer methods utilise cell-level abundance features to identify cell clusters [20], or model distributional differences in expression of individual genes to extract biological meaning of the regulatory network pertinent to cell clusters [21-25]. However, these methods lack more complex gene regulatory information to interpret gene expression networks and are significantly influenced by the data structure [24, 26].

Recent studies such as CADD [27] and UnTANGLeD [28], use statistical methods to quantify probabilities and relationships in the genome to help interpret complex genomic data. These methods are powerful because they provide an independent, unsupervised, simple, and quantifiable analysis of the genome that seamlessly interfaces with orthogonal genomic data to interpret genetic information. We recently demonstrated that histone modification deposition patterns derived from consortium-scale genomic data can be used to discern genes underpinning cell identity [19]. We found that genes frequently harboring broad H3K27me3-domains across diverse cell types significantly enrich for cell-type specific regulatory genes [19]. This approach, which we call TRIAGE (Transcriptional Regulatory Inference Analysis of Gene Expression), is unsupervised and does not depend on external reference data, statistical cutoffs, or prior knowledge. We used consortium data from the Human Cell Atlas, FANTOM, and a draft map of the human proteome to show that TRIAGE provides an independent, orthogonal model to interpret genomic data from any cell or tissue type. TRIAGE devised a genome-wide quantitative feature called repressive tendency score (RTS) which can be used as an independent reference point that infers cell-type regulatory potential for each protein-coding gene.

Here, we present a two-step pipeline drawing on the fundamental principles of TRIAGE to enable analysis of cell clustering and interpretation of cell populations; the approach scales to any cell type and data type, including applications across species to enable analysis of biological diversity in scRNA-seq data.

## Methods

### Identifying priority cell-type-specific regulatory genes

RTS priority genes were obtained by ranking repressive tendency scores (RTS) calculated from EpiMap data across 834 cell types [29] using methods as previously described [19].

### Analysis of RTS priority gene expression correlation with cell differentiation pseudotime

A single-cell RNA sequencing dataset characterising definitive endoderm differentiation using 125 iPSC lines [30] was used to identify genes significantly influencing differentiation potential. The average expression of each RTS priority gene (**Table S1**, genes above the inflection point) in each cell line was calculated at day 0, day 1 and day 2 of endoderm differentiation. A linear regression was performed between the average gene expression at pluripotency and the average pseudotime for each cell line at day 3 of differentiation as calculated in the original data [30]. A false discovery rate correction was applied to account for multiple testing. This analysis was repeated for gene expression at days 1 and 2 of differentiation.

### Pre-Processing of mouse gastrulation atlas data[31]

For the mouse gastrulation atlas data, mouse gene names were converted to human gene names. Genes with no expression in all cells were removed, and low-quality cells were excluded based on *perCellQCMetrics* and *quickPerCellQC* functions from the R *scater* package [32]. Filtered data was normalised by library size calculated from *computeSumFactors* function from *scran* package [33]. Normalised counts were used for *TRIAGE* [19] conversion which generate a discordance score (DS) matrix. A downstream principal component analysis (PCA) was performed using the *RunPCA* function in *Seurat* to compute 50 principal components (PCs), and the 50 PCs were used by uniform manifold approximation and projection (UMAP) through *RunUMAP* function [13].

### Development of TRIAGE-Cluster

#### Density estimation and Peak selection

For each cell in the data set, the RTS priority gene with highest expression for each cell was identified and its corresponding RTS used for downstream analysis as follows: UMAP coordinates were used with the assigned RTS for each cell as weight for estimating kernel density using function *stats*.*gaussian_kde* from *scipy* package [34], which estimates the probability density function of RTS-weighted cell state. For any given cell *x*, the weighted Gaussian kernel density estimation can be calculated as:

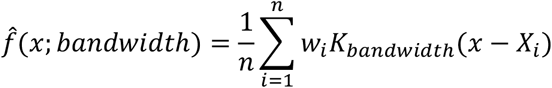

where *n* is the total number of cells, *w*_*i*_ is RTS and *X*_*i*_ the UMAP embeddings of the *i*-th cell, *K* is a Gaussian kernel function. Bandwidth was calculated from Scott’s Rule [35] and was adjusted to suit scRNAseq data shown as following:

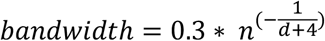

where *n* denotes number of cell and *d* denotes number of dimensions. A density map with contours (contour levels=10) were generated. At each contour level, unsupervised *DBSCAN* from *sklearn* package [36] was used to group cells into spatially separated clusters. From each contour level, regions with local RTS maxima are selected as peaks (i.e., clusters without smaller clusters within them from a higher contour level). Peaks from all contour levels determine the cell cluster diversity from the data set.

#### Jaccard similarity dendrogram

We used Jaccard similarities of each pair of peaks to calculate Euclidean distance matrix. R package *hclust* was used to perform hierarchical clustering and extract the dendrogram representation.

#### Adjusted Rank Index (ARI) analysis

We used *adjustedRandIndex* function from *mclust* R package [37], and ARI was performed using peaks from TRIAGE-Cluster. Original RTS priority gene rank was used as gold standard and accuracy of data structure was evaluated using random removal of genes from RTS priority gene list, removal of RTS priority genes from bottom rank to top, removal of RTS priority genes from top rank to bottom, and random removal of genes from the whole genome. Each removal step is 50 genes.

#### Correlation and Gene Ontology enrichment analysis

We performed Spearman rank correlation analysis using *cor* function from *stats* using whole transcriptome and 1000 highly variable genes. Highly variable genes are defined using *modelGeneVar* and *getTopHVGs* (n=1000) functions from *scran* package. Gene Ontology (GO) enrichment analysis was performed using *topGOtable* function from *pcaExplorer* package with genome wide annotation for human *org*.*Hs*.*eg*.*db*.

### Development of TRIAGE-ParseR

#### Processing of H3K27me3 data

We used consolidated H3K27me3 BED files for 111 Roadmap tissue and cell types [38]. We also downloaded bigwig files representing 834 samples from EpiMap [29] and converted them into *bedgraph* format using *bigWigToBedGraph* [39, 40]. Subsequently, we used *MACS2* [41] to call H3K27me3 enriched loci with - broad option to capture broad deposition of H3K27me3.

#### Extracting H3K27me3 patterns

We performed PCA to extract orthogonal patterns of H3K27me3 depositions from consortium-level epigenomic data [29, 38]. H3K27me3 deposition values were defined by breadth of H3K27me3 peaks overlapping proximal regions of RefSeq annotated transcription start sites (TSS) (+/- 2.5kb) of genes. If multiple peaks overlap a given gene, the broadest peak was chosen. We used top 67 (Roadmap) or 60 (EpiMap) top PCs which explain most of the data variance (i.e., 96.5% or 97.7% respectively). PCA loading values represent unit scale components for covariances between observed H3K27me3 breadths of different samples (**Figure S4a**) while eigen vectors indicate degree of gene’s correlation to H3K27me3 patterns.

#### Analysis of H3K27me3 patterns among top 100 genes ranked by discordance score

We compared underlying H3K27me3 patterns associated with top 100 genes ranked by discordance score which are highly enriched with variably expressed transcription factors between different tissue groups (defined by expression coefficient of variation>1) (**Figure S4h**). To this end, we first analysed enrichment of H3K27me3 patterns across genes. For each gene, empirical probability of observing a given value higher than the 95^th^ percentile of the 26,827 RefSeq genes in each eigen vector was calculated. We define that genes with the empirical p-value of less than 0.05 (right-tail) demonstrate strong association with a given H3K27me3 pattern. To understand whether each of the top 100 genes ranked by discordance score demonstrate significant association with a given pattern, we randomly sampled another 100 genes (null distribution) and counted incidences (*r*) where the observed value is higher than samples in the null distribution. We permuted this process 1,000 times and calculated a probability of gene *i* (*p*_*i*_) to have a value higher than 95% of the random samples.

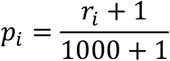

#### Identifying an appropriate cluster number for genes by H3K27me3 patterns

To cluster genes ranked by discordance score based on associated H3K27me3 patterns, we used Bayesian information criterion [42] to find a suitable number of clusters. The BIC finds a suitable number of clusters from distributions of observed H3K27me3 depositions. While we used top 100 genes ranked by discordance score based on their enrichment of regulatory genes (**Figure S4h**), the user can select any meaningful number of genes for this analysis. For a given top 100 TRIAGE-prioritised gene set (i.e., n=100), values of top *m* PCs (i.e., *X*_*i*_ = *x*_*i*,2_, *x*_*i*,1_, …, *x*_*i,m*_, for *i*-th ranked gene where *i* ∈ {1,2, …, 100}) were used as features for the parameter selection. The method then ranks PCs by level of variance observed among the input gene set and selects top *m* PCs (default = 10 PCs). Subsequently, the maximum likelihood to observe PC values from all top 100 genes ranked by discordance score was iteratively calculated with a varying number of clusters *θ* where *θ* ∈ {1,2, …, 100}. The cluster was defined by a Gaussian model *φ*_*j*_(*X*|*μ*_*j*,1_, *μ*_*j*,2_, …, *μ*_*j,m*_, *σ*_*j*,1_, *σ*_*j*,2_, …, *σ*_*j,m*_) for *j-*th cluster centred at means *μ*_*j*_with variances *σ*_*j*_ across *m* PCs, where *m* is 67 or 70 for Roadmap or EpiMap data respectively. The clusters represent a region of high probability mass estimated by expectation-maximisation (EM) algorithm. Intuitively, if *θ* is 1, every gene belongs to the same cluster while if *θ* is 100, every gene belongs to its own cluster. The aim of BIC is to find an appropriate value for *θ* that gives the lowest BIC value (i.e., a cluster number which is the most balanced point between model complexity and likelihood of observing data given the cluster number and model parameters).

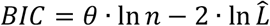

Where 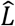 is the maximum value of the likelihood function (L) given the number of clusters (*θ*) and model parameters. The likelihood functions are written as follows.

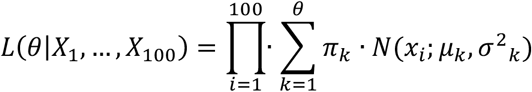

Where *π*_*k*_ represents a probability (or mixture proportion) that gene *X*_*i*_ belonging to *k*-th cluster.

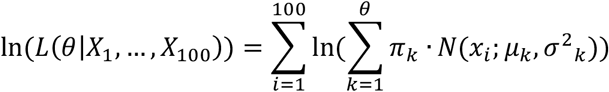

We aim to find *θ* such that the BIC is the smallest. Hence, this problem is equivalent to finding the maximum log-likelihood value given *θ* (i.e., ln 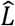).

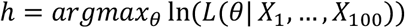

Due to sensitivity of EM method to initial parameter values, this analysis was repeated multiple times (default = 10 times) and the most frequently occurring values were chosen as an appropriate number of clusters. To ensure reproducibility of the results, random state parameter for GaussianMixture function from scikit-learn Python module was set to 42.

#### Assigning genes to H3K27me3 clusters using Gaussian mixture models

After identifying an appropriate number of clusters (*h*) for a given prioritised gene set, the next step is to probabilistically assign each gene into one of the clusters. We assumed multivariate Gaussian distributions for each cluster. The Gaussian Mixture Model (GMM) estimates parameters of the model through the EM algorithm in dimensions defined by selected PCs. Briefly, the EM randomly initiates parameters Θ (i.e., mean and variance of each cluster) then iteratively adjusts these parameters until convergence. By default, iterations stop when log-likelihood gain of the model is less than 1e-3.

For each observation, the expectation step (E) calculates likelihood of the observation belonging to clusters given current model parameters while the maximisation step (M) adjusts these parameters to maximise the likelihood of data. Once the model is trained, posterior probability of the gene to fall into each cluster is calculated. Finally, the gene is assigned to a cluster with the highest posterior probability.

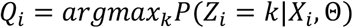

Where *Z*_*i*_ is a latent variable for *i*-th gene being in a *k*-th cluster and *Q*_*i*_ is a cluster where *i*-th gene is finally assigned.

#### Functional annotation of gene clusters

To test functional relevance of gene clustering, protein-protein interaction data as well as GO enrichment analysis using STRING data were performed [43]. Gene clusters with significant PPI (FDR < 0.001) were identified and enriched GO terms were extracted.

#### STRING PPI analysis

To analyse functional connectivity between genes, STRING database was utilised. PPI enrichment was assessed with default settings which include both functional and physical protein associations. Network plots represent both a series of experimental evidence (i.e., protein fusion, neighbourhood, co-occurrence, co-expression and other experimental evidence) and computational predictions (i.e., text mining and database).

### Data visualisation of integrated data in a circular plot

We used *circlize_dendrogram* from the *circlize* R package to curve the Jaccard similarities dendrogram. The *circos-heatmap* function was used to generate the circular plot for the TRIAGE-Cluster peak number, and the proportion of cells in TRIAGE-Cluster peaks based on original annotation and timepoints. The order was defined by the Jaccard similarities dendrogram. All circular plots were compiled in Adobe Illustrator.

Software used in this study are listed in **Table S2.**

## Results

**Figure 1** provides a stepwise schematic for the workflow developed in this study, in which TRIAGE is used to determine cell populations and regulatory gene programs in scRNA-seq data. A glossary of terms is provided in **Table 1**. First, for every protein coding gene, TRIAGE determines the frequency of broad H3K27me3-domains at each locus across 834 EpiMap biosamples. This is referred to as a gene’s repressive tendency score (RTS). Second, any orthogonal input gene expression data set (or any data mapped to protein coding genes) is multiplied with the RTS to derive a discordance score (DS). The DS enables genes with high regulatory potential to be prioritized (ranked by DS value). Third, TRIAGE-Cluster uses the top ranked gene’s RTS value for every cell as a weight to identify cell clusters in a single cell UMAP space. Fourth, we use epigenetic data in a machine learning model to parse pseudo-bulk DS gene lists into biologically functional groups.

**Table 1.** Glossary of terms

|  |  |
| --- | --- |
| RTS | Repressive tendency score. Calculated from broad H3K27me3 domains across 834 cell types. High RTS genes are lowly expressed and have higher cell-type-specific regulatory potential. |
| Discordance score | Discordance score is calculated by multiplying log-transformed normalised gene expression values of any input data set with TRIAGE-derived RTS values for each gene. |
| RTS Priority genes | Genes with RTS above the inflection point of the interpolated RTS curve are used to identify peak regions in the TRIAGE clustering method. |

**Figure 1.**
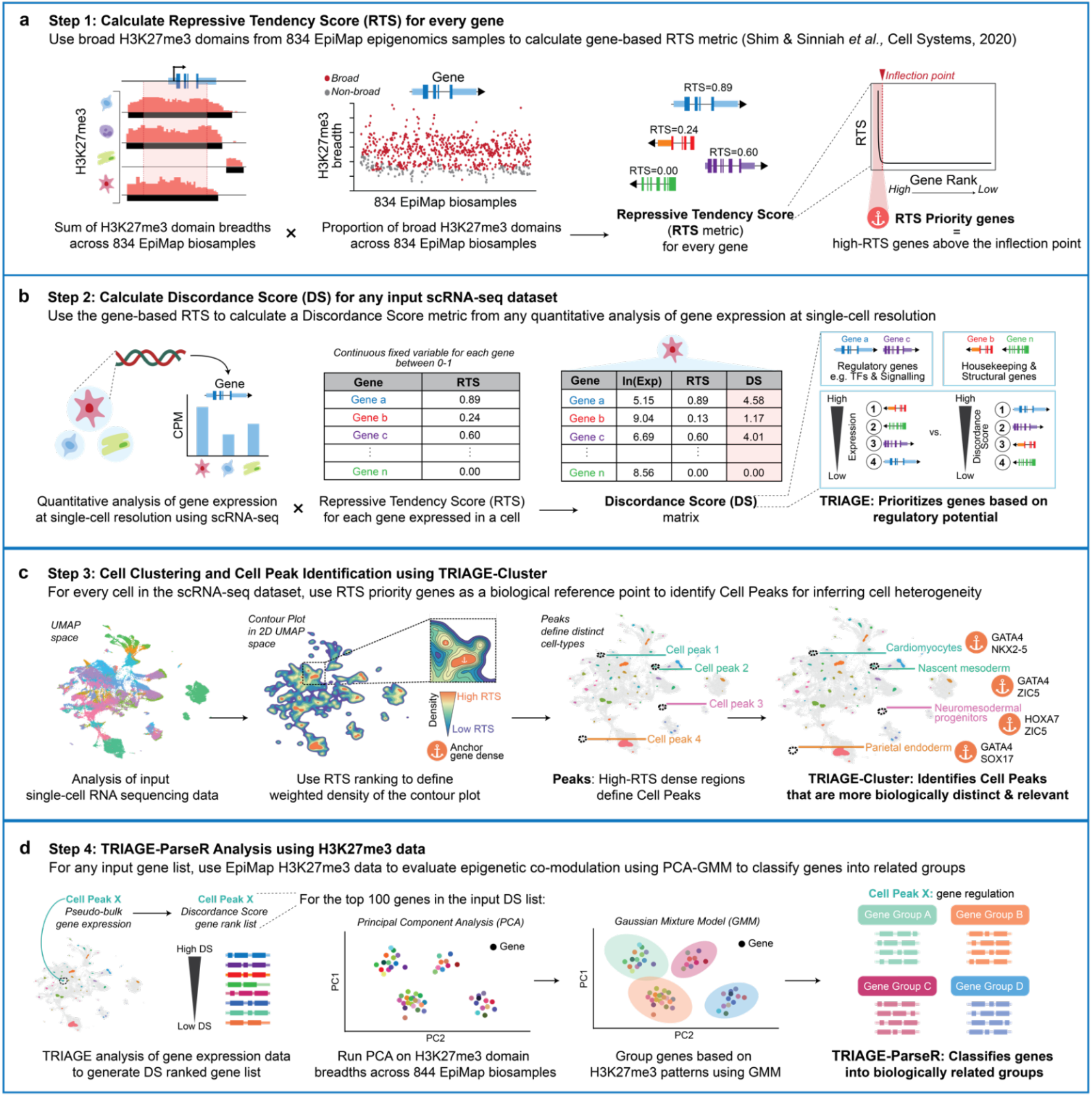
Overview of unsupervised pipeline for analysis of single cell data to identify cell types. **a)** TRIAGE calculates a repressive tendency score (RTS) for every gene based on its association with broad H3K27me3 domains across 834 EpiMap bio-samples [29]. RTS genes above the inflection point of the RTS curve are defined as RTS priority genes to assist in peak identification using TRIAGE-Cluster. **b)** Input single cell expression matrix is transformed to discordance matrix to convert the original expression value to discordance score (DS). The DS results in high ranking of cell type regulatory genes. **c)** We use RTS priority genes in a density-based clustering method, TRIAGE-Cluster, to identify cell populations in UMAP space. **d)** For each peak, genes are ranked by pseudo-bulk discordance score (DS). TRIAGE-ParseR analysis uses PCA and Gaussian mixture model (GMM) to group genes into functional groups to assist with cell type identification.

### Using EpiMap data to calculate genome-wide repressive tendency scores

While our first study [19] developed RTS using 111 NIH Epigenome Roadmap H3K27me3 samples, this study expands the reference data for calculating RTS by measuring H3K27me3 deposition across 834 cell samples and 27 tissue groups in EpiMap [29] (**Tables S3**). We identify 993 priority genes above the inflection point of the interpolated RTS curve (RTS > 0.013) (**Figure 2a**). Highly variably expressed transcription factors (TFs) represent a positive gene set of cell type specific regulatory genes [19] and are significantly enriched among RTS priority genes (**Figure 2b**). Furthermore, genes ranked highly by RTS tend to be more cell type specific and lowly expressed (**Figure 2c-e**), a profile typical of regulatory factors, like TFs.

**Figure 2.**
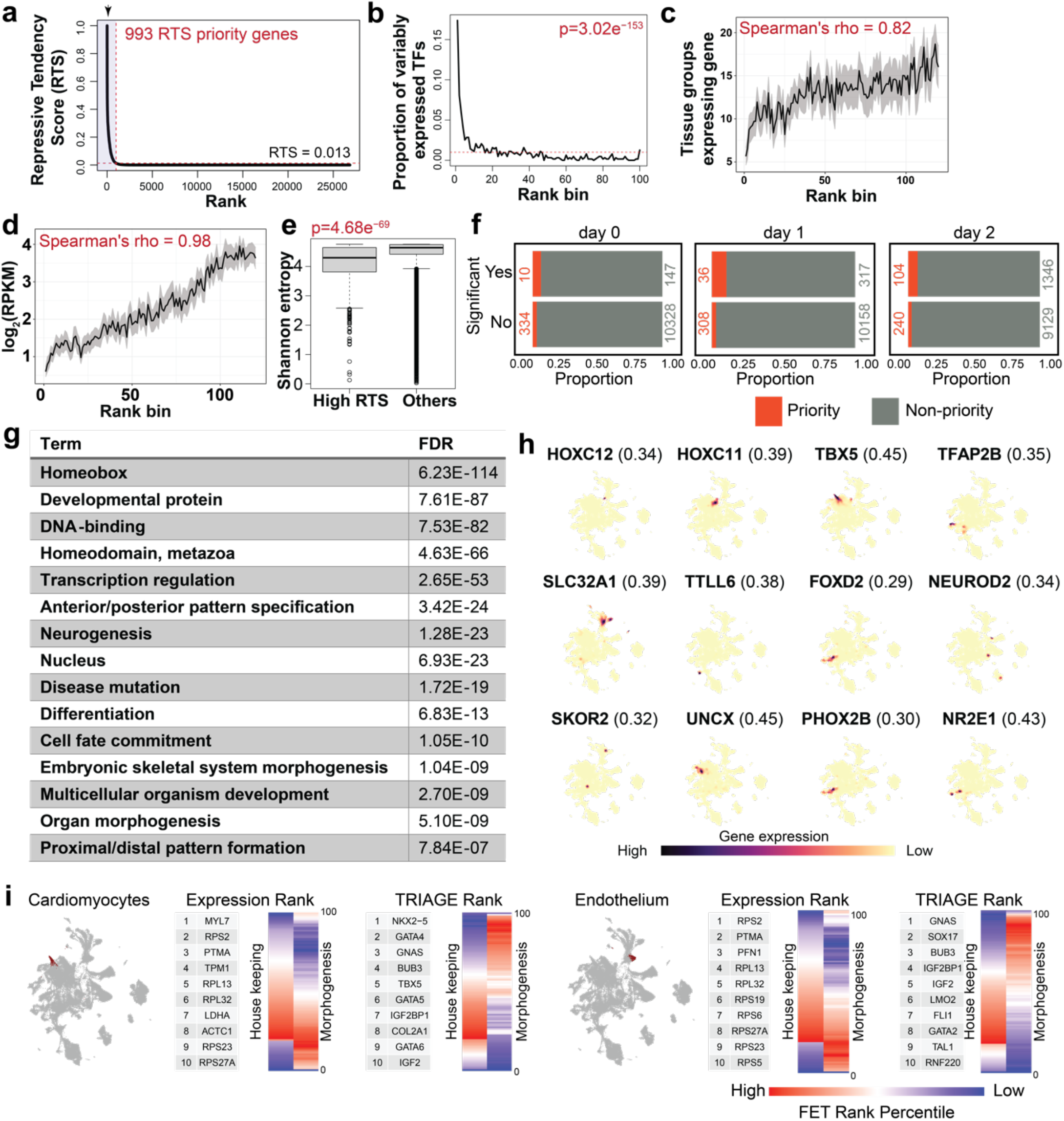
TRIAGE is a gene quantification and selection strategy providing a biological reference point for cell-specific genetic features. **a)** 993 genes (RTS priority genes) have RTS values above the inflection point of the interpolated curve (red dashed line shows RTS = 0.013). **b)** High-RTS genes are significantly associated with variably expressed transcription factors (VETFs). One-tailed Fisher’s exact test shows the top 5%, representing 1,340 RTS genes enriched with VETFs (p = 3.023e^-153^). Red dashed line represents the uniform distribution (proportion = 0.01). VETFs comprise 634 transcription factors (TFs) with variable expression across 46 Roadmap cell types [19, 48]. **c-e)** Genes with high-RTS are associated with cell/tissue specificity **(c)**, are lowly expressed **(d)** and have higher entropy **(e).** **f)** Odds Ratio (OR) analysis demonstrating that RTS priority genes significantly influence the pseudotime trajectory of cell differentiation based on analysis of single-cell sequencing data characterizing definitive endoderm differentiation from 125 iPSC cell lines [30]. The actual number of genes for each section are listed inside each plot. **g)** Gene Ontology terms enriched in the top 993 RTS priority genes are significantly enriched in gene programs controlling cell identity, differentiation, and development. **h)** UMAP plots of mouse gastrulation [31] showing selected RTS priority genes in the UMAP space indicative of cell-specific expression. RTS priority genes and their respective RTS are shown in parentheses. Mouse gene names were converted to human gene names for TRIAGE transformation. **i)** TRIAGE transformation of gene expression re-ranks genes to enrich for genes with regulatory potential. This is shown for pseudo-bulk gene expression analysis of annotated cardiomyocytes (left) and endothelial cells (right) from [31]. Data show UMAP of annotated cell types (left) followed by comparison of original expression vs TRIAGE gene ranking using top 10 ranked genes. The heatmaps showing distribution of housekeeping and morphogenesis genes ranked by original expression and TRIAGE. Enrichment is calculated for each rank bin relative to all genes (one-tailed Fisher’s exact test).

We evaluated whether the expression level of RTS priority genes influences cell differentiation more than would be expected by random chance. To test this, we analysed expression of RTS prioritised genes to evaluate their expression abundance correlated with differentiation trajectory using pseudotime analysis of definitive endoderm over three days of differentiation measured from 125 iPSC lines [30]. These data show that expression of RTS priority genes between day 0-2 significantly affects the differentiation potential of cells (Odds Ratio (OR) on day 0 = 2.10, p = 0.04, day 1 = 3.75, p = 7×10^−14^, and day 2 = 2.94, p = 2.7×10^−20^) (**Figure 2f**). This is borne out by data showing that high-RTS genes are enriched in DNA-binding factors such as homeobox genes (FDR 6.2e^-114^) controlling fundamental processes in cell differentiation, development, and tissue morphogenesis (**Figure 2g)**.

We next tested whether the RTS values could be used as an orthogonal metric to identify cell types in scRNA-seq data. We evaluated data sets for method development and selected an atlas of mouse embryonic development which profiles 116,312 cells collected at nine sequential time-points ranging from e6.5 to e8.5 days post-fertilisation [31] (**Figure S1, S2** and **Table S3**). These data comprise all primary germ layers, consisting of 37 classified cell types including ectoderm, mesoderm, and endoderm (**Figure S2**). We first find that high-RTS ranked genes have cell-specific expression patterns across the UMAP space (**Figure 2h**). Second, analysis of pseudo-bulk gene expression of two annotated cell types, cardiomyocytes and endothelial cells shows that original gene expression enriches for housekeeping and functional genes, while the TRIAGE transformation to DS enriches for regulatory genes underpinning the mechanistic basis of heart (*NKX2-5, GATA4, TBX5*) and endothelial (*SOX17, GATA2, TAL1, LMO2*) development and morphogenesis (**Figure 2i**).

### TRIAGE RTS demarcates cell specific clusters in scRNA-seq data

**Figure 3a** demonstrates how RTS assigned to genes by TRIAGE quantifies a distinction between non-specific vs cell type specific genes. For example, *EOMES* represents a broad mesendoderm cell type (RTS 0.19), while more specific markers of lateral plate mesoderm (*HAND1*, RTS: 0.28) and cardiomyocytes (*NKX2-5*, RTS: 0.35) demonstrate that RTS values increase according to cell specificity. Similar examples are shown for paraxial mesoderm and endoderm (**Figure 3a**).

**Figure 3.**
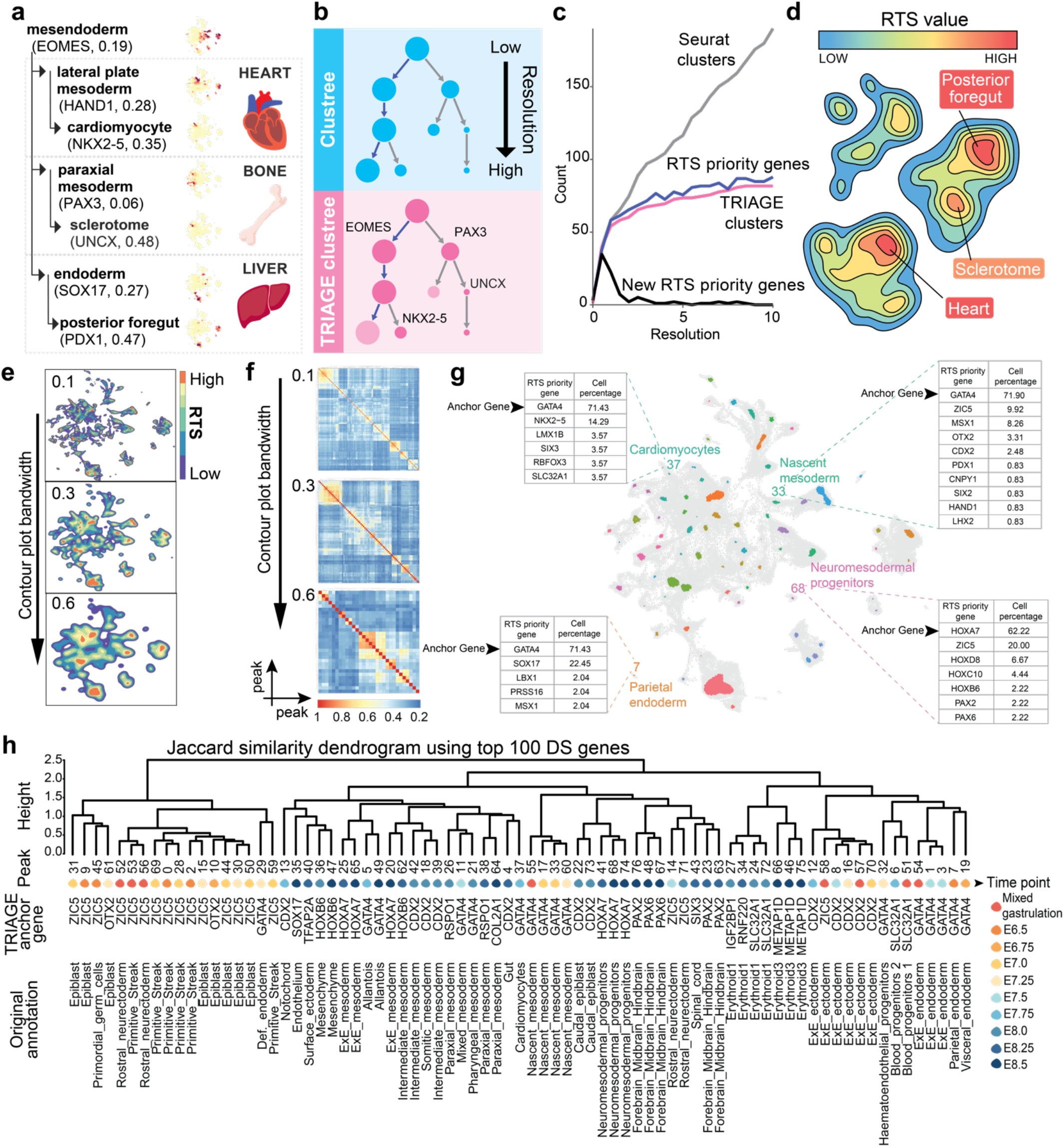
TRIAGE-Cluster is an unsupervised approach to enable cell type identification in a UMAP space. **a-b)** Example **(a)** and schematic **(b)** demonstrating how RTS priority genes provide a quantitative value assigned to all genes which can be exploited for cell-type identification. **(a)** The quantitative value of RTS priority genes enables identification of specific cell types where increasing RTS values correlate with increasing cell type specificity. UMAPs in the middle show expression of selected priority genes. **(b)** Strategy for implementing decision making criteria when splitting clusters from low to high resolution in normal clustering and TRIAGE-Cluster. **c)** Evaluation of TRIAGE decision-making criteria using Seurat clustering resolutions. Seurat cluster: number of clusters at each Seurat resolution. RTS priority genes: cumulative number of RTS priority genes across each Seurat resolution. TRIAGE clusters: number of clusters with at least one RTS priority gene expressed at each Seurat resolution. New RTS priority gene: number of new RTS priority genes at each resolution. **d)** Schematic concept for using RTS priority genes as biological anchoring approach to analyse single cell data sets. We use a weighted density analysis of RTS priority genes in each cell to derive a topographic landscape view of the data with defined cell types identified as peaks in the contour plot. **e)** Contour plot analysis applied to single cell atlas of mouse gastrulation [31] with granularity from high to low in visualisation using parameter bandwidth setting. **f)** Heatmaps with bandwidth resolutions (**panel e)** showing pairwise Jaccard similarity score between peaks calculated from gene discordance score for each cluster. **g)** UMAP of single cell atlas of mouse gastrulation using TRIAGE-Cluster with bandwidth setting of 0.3. Four examples show the RTS priority genes (anchor genes) expressed in the majority of cells within the peak and annotation assigned by the original classification [31]. **h)** Jaccard similarity dendrogram with bandwidth 0.3 (**panel f**) showing peak relationship (first row), mapped timepoint (second row), anchor gene (third row), and mapped original annotation (forth row).

We hypothesised that RTS assigned to genes could provide a method to identify gene clusters in single cell data. We designed a proof-of-concept analysis comparing Seurat clustering vs RTS priority genes as a decision-making feature influencing clustering tree analysis over 10 resolutions (**Figure 3b**). Standard Seurat clustering progressively divides clusters with increasing resolution (**Figure 3c**). For TRIAGE-Cluster, we assumed enrichment of RTS priority genes in a cell population demarcates unique cell types and evaluated this at each stepwise resolution of clustering. We found that RTS priority genes characterising new clusters quickly plateaus demonstrating that increased clustering resolutions do not necessarily imply biologically distinct cell types. Furthermore, RTS priority genes trimmed the number of clusters, while still being sensitive enough to identify new clusters even at very high resolutions, representing rare cell types that cannot be detected in broad categorizations at low resolutions (**Figure 3c**).

### Weighted density analysis of RTS priority genes determines cell clustering

Based on these observations, we designed an approach drawing on the concept of a topographic map where contour lines demarcate peaks of cell specificity in the UMAP space (**Figure 3d**). We hypothesised that RTS prioritised genes expressed in cells demarcate cell-specific identity and a gene’s RTS can define contour lines demarcating peaks (specific cell types) from valleys (transitional or non-specific cell types) (**Figure 3d**). Using weighted density estimation, the RTS is used as the weighting parameter to adjust cell density in a whole transcriptome derived UMAP space.

We applied this method to the mouse gastrulation atlas data [31] (**Figure 3e-h**). Bandwidth resolutions are shown for the contour plot (**Figure 3e**) and pairwise Jaccard similarity between peaks (**Figure 3f**). **Figure 3g** shows the cell clustering result at bandwidth 0.3 identifies 77 peaks. For selected populations we show how high-RTS genes, which we call anchor genes, reveal identity-defining genes across different cell types in the data. For example, cardiomyocytes are anchored primarily by the mes-endoderm-associated gene *GATA4* and cardiac regulatory factor *NKX2-5*. Parietal endoderm is anchored by *GATA4* and the endoderm-associated transcription factor *SOX17*.

We next evaluated cell-cell relationships derived by k-means clustering of the pairwise Jaccard similarity analysis of the mouse gastrulating embryo (**Figure 3f**). The output provides an integrated view of TRIAGE-Cluster derived cell relationships in which we show 1) the output dendrogram organising cell relationships using each cluster’s DS, 2) each cluster’s time point as it appears during embryonic development, 3) the clusters original annotation from the mouse gastrulation atlas and 4) the highest expressing RTS priority gene which we call the cluster’s “anchor gene” (**Figure 3h** and **Figure S3**).

### TRIAGE-Cluster captures granular cell diversity in scRNA-seq data

We next analysed the sensitivity of TRIAGE-Cluster to parameter settings that help determine the clusters. We first asked whether removal of RTS priority genes influences the clustering output using an adjusted rand index (ARI) analysis. While random removal of genes from the genome has no impact on TRIAGE-Cluster accuracy, it is not unexpected that the most dramatic impact on clustering accuracy occurs with removal of high-RTS genes (**Figure 4a**). This is further demonstrated across thirteen independent *in vivo* and *in vitro* scRNA-seq data sets which shows that high-RTS genes are the primary drivers of clustering across all data sets (**Figure 4b**).

**Figure 4.**
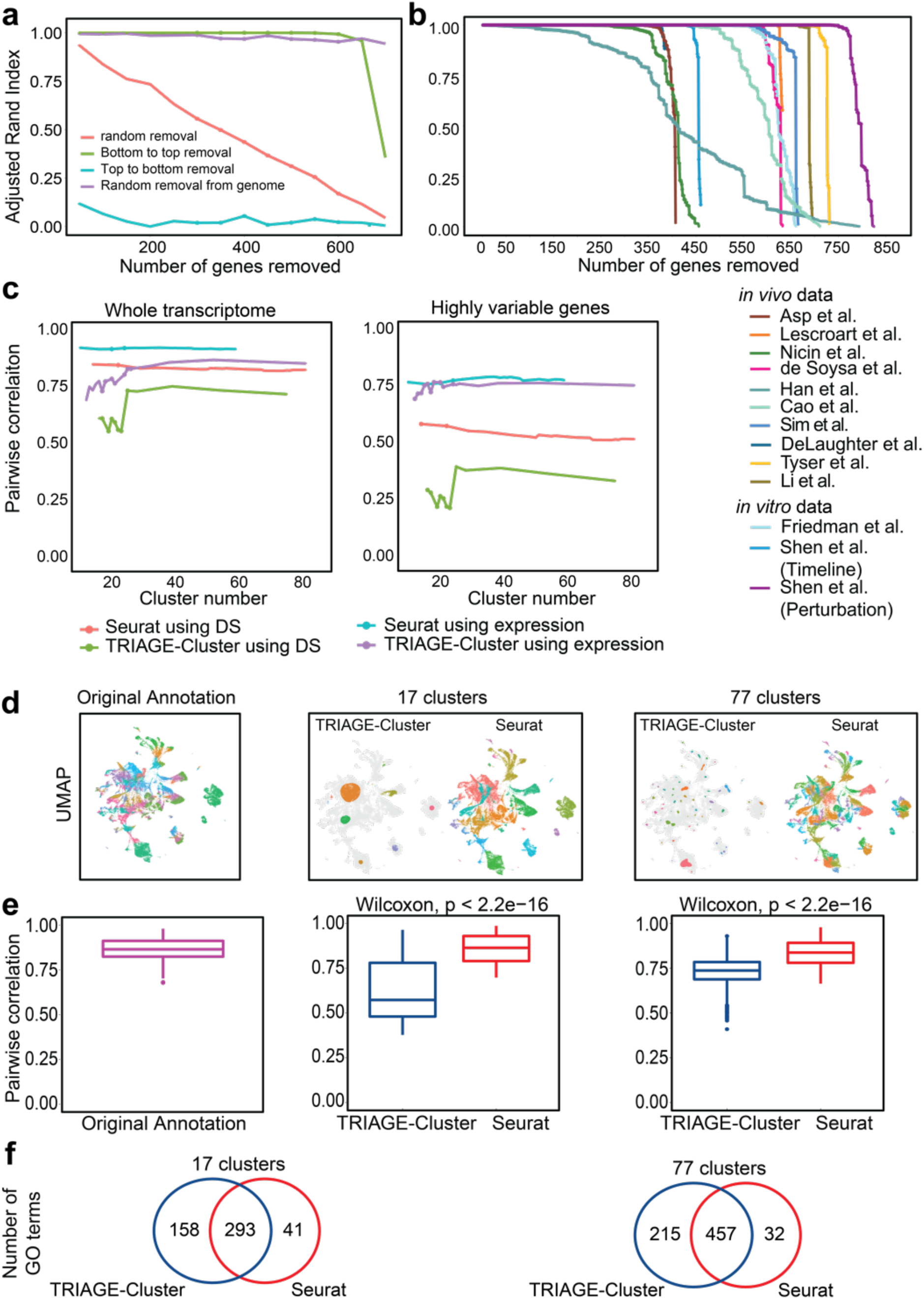
TRIAGE-Cluster identifies biologically distinct cell types. **a)** Adjusted Rand Index analysis of single cell atlas of mouse gastrulation [31] demonstrating that the quantitative RTS priority gene rank is required for accurate cell clustering. **b)** Thirteen data sets from *in vivo* and *in vitro* cell atlases analysed for clustering performance by removal of RTS priority genes from bottom to top demonstrate a critical role for high-RTS genes in clustering accuracy. **c)** Comparison of peak-peak similarity using Spearman rank correlation of whole transcriptome (left) and highly variable genes (right). We evaluate the correlation with the mouse gastrulation atlas data [31] of original expression and TRIAGE transformed matrix (DS matrix), showing Seurat and TRIAGE-Cluster with increased number of clusters at 20 resolution/bandwidths (0.1-2 with 0.1 increments). **d)** UMAPs of the mouse gastrulation atlas data showing TRIAGE-Cluster and Seurat peaks. Quantitative analysis demonstrates peak-peak correlation comparing original annotation cell diversity compared to TRIAGE-Cluster and Seurat when cell diversity is fixed at 17 or 77 clusters. **e-f)** Spearman rank correlation **(e)** and Gene Ontology analysis **(f)** demonstrate that TRIAGE-Cluster gives significantly lower correlation and therefore captures greater differences in cell diversity **(e)** and supported by increased gene ontology enrichment compared to Seurat or original annotations (**f**).

We next tested the performance of TRIAGE-Cluster using the assumption that distinct cell types have gene expression differences reflected by lower cluster-cluster pairwise correlation. Comparing TRIAGE-Cluster against Seurat, we tested gene expression similarity between all pairwise peaks at different clustering resolutions using Spearman rank correlation. Seurat clustering shows highly stable similarity between cell types at all clustering resolutions regardless of input data type (original expression or DS, i.e., when expression is multiplied by RTS). Furthermore, Seurat shows stable correlation between clusters using either the whole transcriptome or highly variable genes for gene expression similarity comparison. This suggests that separation of distinct cell types over different cluster resolutions in Seurat is not driven by significant global gene expression differences (**Figure 4c**).

In contrast, TRIAGE-Cluster peaks consistently result in lower peak-peak Spearman rank correlation compared to Seurat and the correlation increases at higher clustering resolutions as differences between cell types become less distinct (**Figure 4c**). We achieved the lowest peak-peak correlation using TRIAGE-Cluster with highly variable DS values assigned to genes. We further demonstrate this comparing peak-peak correlations of TRIAGE-Cluster vs Seurat clusters in which the clustering resolution of both methods results in the same cluster number (**Figure 4d)**. Peak-peak correlations of cell types originally defined in the single cell atlas of mouse gastrulation [31]were similar to those defined by Seurat (**Figure 4d-e)**. To further validate that TRIAGE-Cluster peaks capture functionally relevant gene sets in the data, we performed Gene Ontology analysis using the genes with the top 100 DS for each peak or cluster. These data show that TRIAGE-Cluster peaks have more GO BP terms than Seurat clusters and therefore demonstrates that TRIAGE-Cluster extracts genes involved in more diverse biological processes (**Figure 4f, Table S4-7**). Taken together, these data show that a combination of DS score with TRIAGE-Cluster efficiently identifies biologically distinct cell types in scRNA-seq data.

In our companion study, we demonstrate the utility of TRIAGE-Cluster by applying it on a multiplexed single cell atlas of iPSC differentiation data that integrates temporal data across eight time points and signalling data with nine developmental signalling perturbation [44]. 48 peaks were identified from the *in vitro* differentiation atlas data, and clustering performance was evaluated using a trajectory approach VIA which evaluates cell states relationship [45]. Comparing analysis using all cells vs TRIAGE-Cluster peaks, TRIAGE-Cluster peaks give better peak-to-node assignment in the trajectory and significant correlation between pseudotime and actual time. These data demonstrate that TRIAGE-Cluster can define the biological diversity and improve cell-cell trajectory analysis in scRNA-seq data.

### TRIAGE-ParseR enhances identification of cell gene regulatory networks

To complement TRIAGE-Cluster, we developed a method called TRIAGE-ParseR to extract regulatory information from a distinct population of cells. We aimed to develop a method that distinguishes biologically meaningful gene groups from gene expression without reliance on subjective external reference points (such as differential expression) or prior knowledge (such as GO, STRING, marker gene selection, or gene class biases) [18].

Based on the rationale that expression of key developmental genes is modulated by both gain or loss of H3K27me3 (Boyer, 2006; Mikkelsen, 2007), we hypothesized that principal component analysis (PCA) of (breadth of) H3K27me3 for genes across diverse cell states, specifically in diverse human tissue and cell types, could help separate functionally relevant gene-gene relationships. After performing PCA, we evaluated the input genes list for shared H3K27me3 deposition patterns encoded by the top principal components (PCs) and modelled by a Gaussian mixture model (GMM). To generate the input gene list, the original expression matrix of a cell type is converted to discordance score and evaluated using TRIAGE-ParseR (**Figure 5a**).

**Figure 5.**
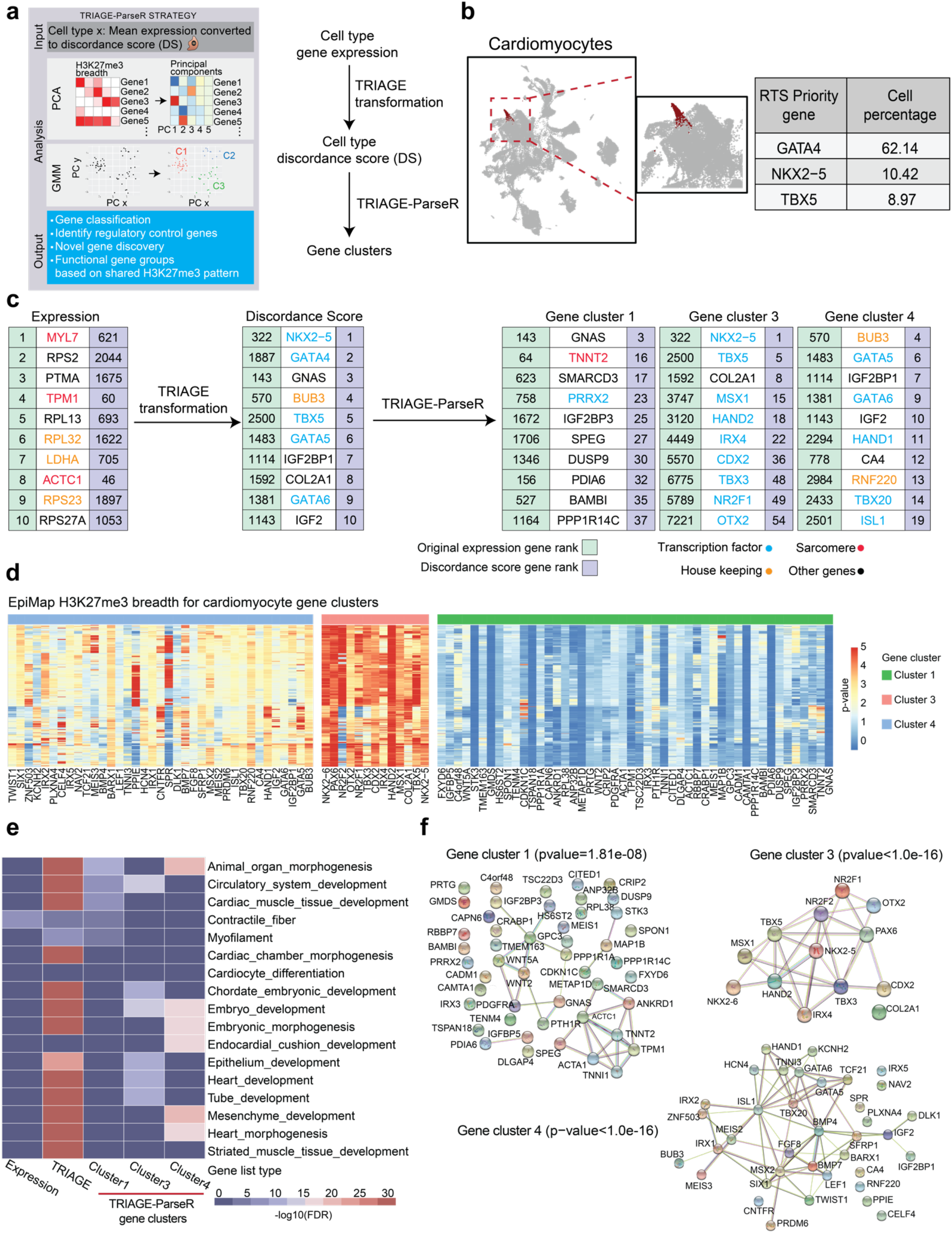
TRIAGE-ParseR clusters genes into groups using unsupervised analysis of epigenetic data. **a)** Schematic workflow describing TRIAGE-ParseR which uses H3K27me3 across hundreds of bio-samples to cluster genes using GMM on PCA space to improve analysis of cell gene regulatory networks. **b)** UMAP of cardiomyocytes from mouse gastrulation atlas data showing top 3 RTS priority genes expressed in cardiomyocytes. **c)** Top 10 genes in annotated cardiomyocytes ranked by original expression (left), discordance score (middle), and top 10 genes in gene clusters resulting from TRIAGE-ParseR. **d)** Abundance of H3K27me3 deposited across each gene locus measured from EpiMap bio-samples demonstrate unique epigenetic patterns demarcating specific gene groups shown in panel c. **e)** Heatmap showing gene ontology enrichment (-log10 (FDR)) across top 100 ranked genes from cardiomyocytes ranked by original expression (first column), discordance score (second column), and TRIAGE-ParseR gene groups (last three columns). **f)** String gene network analysis showing protein-protein interactions of the three cardiomyocyte gene clusters.

Extracted H3K27me3 patterns by PCA showed that (i) most of the data variance can be explained by a small number of PCs (**Figure S4a**), [34] distinct tissue types can be characterised by these H3K27me3 patterns (**Figure S4c**), and genes enriched with a specific H3K27me3 pattern demonstrate tissue-specific biological processes (**Figure S4b**). Furthermore, these data show that the first PC recovers genes with strong tendency to deposit broad H3K27me3 across many tissue types (i.e., high-RTS genes) while subsequent PCs capture more tissue type-specific patterns (**Figure S4d**). This justifies the importance of incorporating diverse PCs to reveal biologically informative signals that distinguish gene programs underpinning different cell types. We used TRIAGE-ParseR to evaluate gene expression from heart tissue and blood (**Figure S4e-f**). In addition to the first PC which shows high concordance with RTS, analysis of diverse PCs reveals unique gene-gene TRIAGE-ParseR signatures that can be used to inform gene-gene relationships.

We applied TRIAGE-ParseR on single cell clusters identified in the mouse gastrulation atlas and focussed the initial demonstration of the method on cardiomyocytes (**Figure 5b**). In the workflow pipeline (**Figure 5c**), pseudo-bulk gene expression (top ranked are primarily housekeeping and structural genes) is transformed with TRIAGE to derive the DS (top ranked are primarily cardiac regulatory genes). The genes with the top 100 DS are used as input to TRIAGE-ParseR (**Figure 5c**) resulting in three gene clusters with significant enrichment by STRING analysis (protein-protein interaction FDR<0.001). **Figure 5d** shows the raw H3K27me3 deposition patterns that clearly demarcate common epigenetic patterns governing each gene group.

We compared GO enrichment at each step of the workflow (**Figure 5c**). While original gene expression shows enrichment in terms related to structural features of cardiomyocytes, TRIAGE discordance score provides robust enrichment in diverse pathways associated with cardiomyocyte morphogenesis and differentiation, TRIAGE-ParseR separates these genes into groups. Genes in cluster 3 (the most significantly repressed genes across diverse cell types, **Figure 5d**) are enriched with heart development terms including diverse transcription factors known to be critical for heart development (**Figure 5e**). Clusters 4 genes have intermediate repression (**Figure 5d**) and are enriched in generic genes governing embryo development (**Figure 5e**). Lastly, genes in cluster 1 have minimal repression across diverse cell types (**Figure 5d**) and are enriched in genes involved in cardiac muscle function (**Figure 5e**). Indeed, STRING analysis revealed that genes in these clusters formed significant protein-protein interactions and therefore revealed both known and unknown gene-gene relationships (PPIs, enrichment p<0.01) (**Figure 5f**). These data demonstrate that H3K27me3 patterns reveal gene regulatory networks relevant to biological functions of genes and TRIAGE-ParseR can meaningfully segregate genes to facilitate cell type identification and biological interpretation.

### Integrating TRIAGE-Cluster and TRIAGE-ParseR to study single cell data

We integrated TRIAGE-Cluster and TRIAGE-ParseR to evaluate cell types in the mouse gastrulation atlas. **Figure 6a** shows a holistic view of all TRIAGE-Cluster peaks, peak relationship at bandwidth 0.3, as well as corresponding original annotation and timepoint. While the mouse gastrulation data set provides a powerful resource for studying the earliest stages of mammalian organogenesis, most cell classifications capture redundant or broad cell groups that lack the nuance of cell diversification during early development. We therefore implement TRIAGE-Cluster as a complementary strategy for identifying more distinct cell types based on its ability to determine local maxima of RTS-ranked genes in the data set as a surrogate indicator of biologically distinct cell populations.

**Figure 6.**
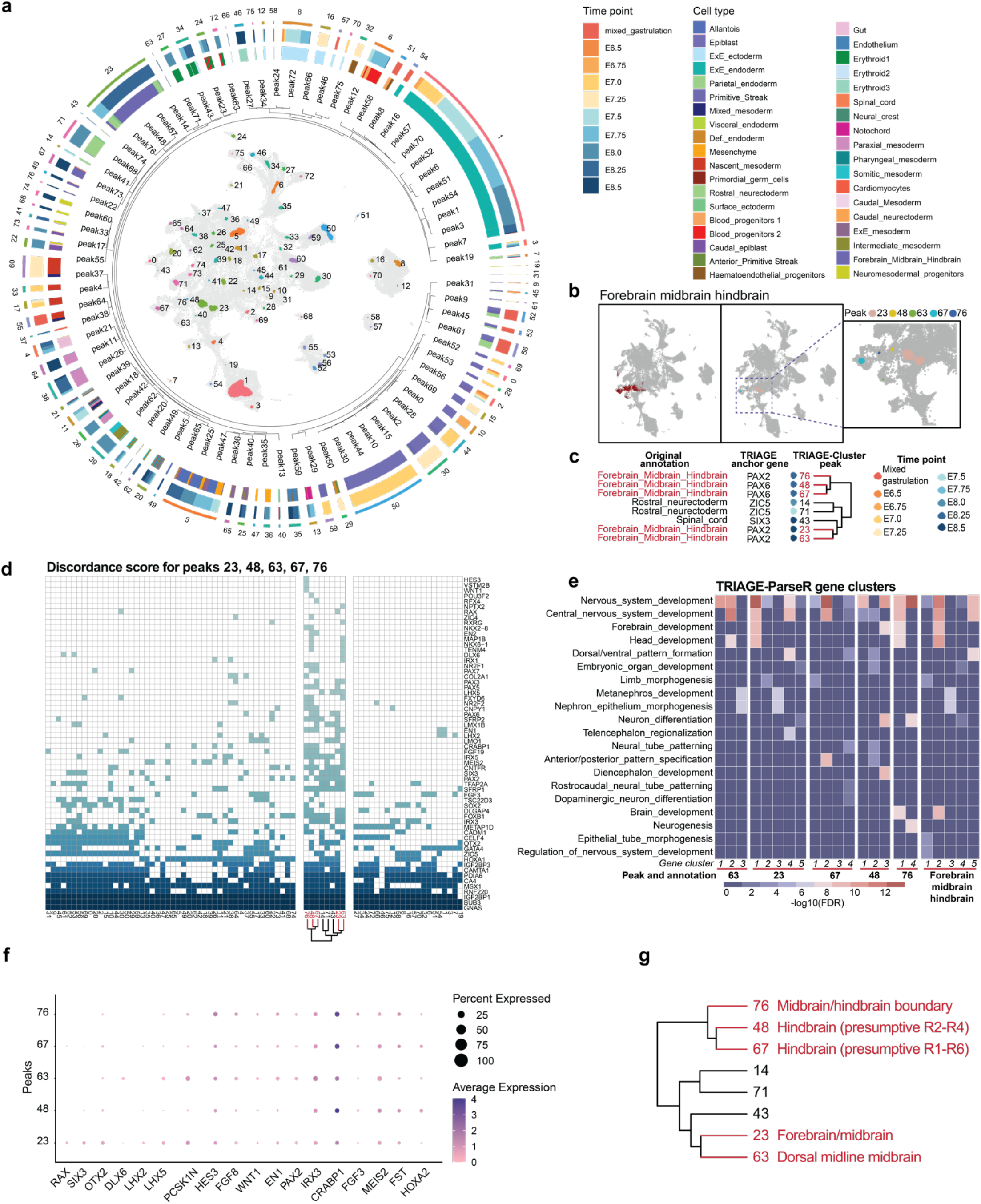
TRIAGE-Cluster enables efficient and unsupervised identification of cell subtype diversity in single-cell data. **a)** Integrated metadata for all peaks identified by TRIAGE-Cluster analysis in the mouse gastrulation atlas data. From inside to out: data include UMAP identifying TRIAGE-Cluster peaks, peak relationships based on 0.3 bandwidth dendrogram (**Figure 3h**), TRIAGE-Cluster peak number, proportion of cells in TRIAGE-Cluster peaks based on original annotation and timepoints. **b)** UMAP of original annotation of forebrain midbrain hindbrain (left) and location of TRIAGE-Cluster peaks (right and inset). **c)** Jaccard similarity dendrogram with bandwidth 0.3 showing five distinct “forebrain midbrain hindbrain” TRIAGE-Cluster peaks with their respective mapped timepoint and anchor genes (extracted from **Figure 2h**). **d)** Specificity of discordance score genes identified for the five “forebrain midbrain hindbrain” peaks compared to all TRIAGE-Cluster peaks. A coloured box indicates a gene (row) characterises that peak (column). Peaks sharing a characteristic gene indicates shared regulatory or structural features defining that cell type. **e)** Heatmap showing different neural cell subtype gene ontology enrichment (-log10 (FDR)) from TRIAGE-ParseR across gene clusters in the five forebrain midbrain hindbrain peaks (the first five blocks) and original forebrain midbrain hindbrain cells (the last block). **f)** Dot plot showing marker genes associated with neural lineage subtypes in each forebrain midbrain hindbrain peak. **g)** Jaccard similarity dendrogram showing revised cell group classifications for the original forebrain_midbrain_hindbrain cell type using integrated analysis of TRIAGE-Cluster and TRIAGE-ParseR.

We selected cells annotated as forebrain/midbrain/hindbrain in the original data. TRIAGE analysis identifies five peaks (23, 48, 63, 67, and 76) mapped to this annotation **(Figure 6b)**. We provide a multi-scale data output to evaluate identity-defining features of each cell type including their relationships defined by TRIAGE-derived dendrogram (**Figure 6c**), the highest ranked DS genes for each cell type **(Figure 6d)**, and TRIAGE-ParseR analysis of gene-gene groupings analysed by GO analysis **(Figure 6e)**. These data reveal a separation of distinct cell relationships within this broad cell grouping that suggest more nuanced cell subtypes governing ectoderm development. Clusters 48 and 67 share high degrees of similarity based on their gene ontology enrichment for neural patterning (Neural tube, Dorsal/Ventral and Anterior/posterior pattern specification, **Figure 6e)** and DS with genes including the shared TRIAGE anchor gene *PAX6*, an organizer of hindbrain patterning. Detailed gene expression data suggest cluster 67 is boundary-defining cells of presumptive rhombomeres R1-R6 (*CRABP1* and DS for *FGF3*) while cluster 48 is more specifically associated with presumptive R2-R4 (*FST* and *HOXA2*, **Figure 6f)**. Cluster 76, anchored by *PAX2*, shares a broad range of identity defining genes related to establishment of the Isthmic organizer at the midbrain/hindbrain boundary (*HES3, FGF8, WNT1, EN1*). In contrast, clusters 23 and 63 share similarity based on the TRIAGE-dendrogram (**Figure 6c**). Cluster 23 shows signatures of Forebrain/midbrain marked by *RAX, SIX3, LHX2, LHX5*, and *PCSK1N* **(Figure 6f)** and GO enrichment for Forebrain development **(Figure 6e)**. Cluster 63 is associated with the dorsal midline of the midbrain marked by *PCSK1N, HES3, FGF8* and *DLX6*. We provide a second analysis of paraxial mesoderm lineage cell types based on detailed dissection using TRIAGE-associated outputs (**Figure S5**). Further examples of the data interpretation pipeline outlined here are provided for analysis of *in vitro* differentiation from pluripotency outlined in the companion paper [44]. The collective results of both studies including new data, software package downloads, and data analysis dashboard can be accessed online (http://cellfateexplorer.d24h.hk/).

## Discussion

Understanding biological relationships between cell types and the gene networks that govern these differences remains central to maximising the value of technologies capturing genomic data at single cell resolution. This study demonstrates that tools like TRIAGE that use an orthogonal reference point to determine identity-defining features provide a powerful mechanism for defining and interpreting the genetic basis of cell populations. Values assigned to all genes based on their RTS can act as an independent biological reference point implemented downstream of dimensionality reduction algorithms to cluster cell types.

Regulation of the genome is in part mediated through histone modifications that control genome accessibility, acting as an on/off switch to regulate gene programs [46]. We reasoned that regions with the broadest histone modification domains (representing extreme levels of on/off control) could be used to quantify the probability that a locus has a role controlling cell decisions and functions. Based on this rationale, TRIAGE [19] uses patterns of histone methylation data to identify regions of the genome that are enriched in cell-specific identity defining genes. Genes with a high RTS have consistent repressive chromatin in most cell types and, therefore, in the rare instances when these genes get turned on, they likely have key roles controlling cell decisions or functions.

The collective suite of methods built around TRIAGE provide several major advantages: 1) TRIAGE is a customisable metric with a single value assigned to genes which makes it simple to implement in any genomic data set, 2) it is versatile, providing a quantitative genomic model to weight gene prioritisation using routine genetic analysis tools, 3) the simplicity of TRIAGE enables broad implementation in data analysis pipelines for any data sample from any cell, tissue, disease, or individual and providing a method to enrich for factors most likely to cause phenotypic changes in cell identity, and 4) can help prioritise genetic targets for drug development and identifying the primary determinants of disease and development.

TRIAGE-ParseR provides several critical advances in studying gene-gene relationships from input expression data. Namely, 1) it provides an unsupervised approach to study the gene regulatory basis of cells without drawing on external reference points (like differential expression analysis) or prior knowledge, 2) it parses genes into groups to reveal biological mechanisms underpinning cell states, and 3) uses an independent reference data set for evaluating epigenetic patters which provides a mechanism for applying this method to any input gene list.

In future studies we aim to address three limitations in the current workflow. First, we demonstrate that the current clustering method significantly restricts cell selection to capture clusters with the most biologically distinct identity. While this is a major advantage, it comes at the cost of data trimming which should be weighed relative to the loss of statistical power provided by the scale of cell sampling. To address this limitation, we aim to use TRIAGE-Cluster in combination with data imputation or trajectory inference methods to interpret cell-cell relationships using all cells in the data set. Second, the current method is reliant on assumptions built into dimensionality reduction representing cell relationships in UMAP space. While useful, major limitations exist for simplifying the data into these graphical presentations. To address this, we propose additional strategies such as comparison of cell types using TRIAGE-dendrograms (**Figure 3h**) which provide independent approaches to evaluate cell-cell relationships. We aim to develop approaches that provide the analysis value of TRIAGE-Cluster but avoid the limitations associated with topographical map representations. Third, TRIAGE-ParseR provides a unique approach to decipher gene-gene relationships but draws on only a single epigenetic reference point. Future developments of this method need to parameter test additional histone modifications representing active promoters (H3K4me3) or enhancers (H3K27ac and H3K4me1) as complementary to repressive chromatin marked by H3K27me3 as a basis for more nuanced analysis of gene-gene relationships.

This study provides a unique strategy for cell type identification using epigenetic information as biological reference point for cell clustering and deconstructing gene modules in single cell expression data. We provide examples of the workflow by analysing *in vivo* organogenesis and *in vitro* differentiation atlases. Collectively, TRIAGE highly ranks regions of the genome enriched in genetic features with profound effects on cell identity because these regions encode potent determinants of cell developmental processes. These ranking features and genome-wide predictions make TRIAGE unique among genomic analysis methods and position it to address areas of need in cell biology and genetics. In addition to our prior studies [19], TRIAGE-Cluster and TRIAGE-ParseR provide a growing portfolio of methods, complementing baseline methods like Seurat, to facilitate analysis of large-scale data sets by integrating genome-wide epigenetic data with scRNA-seq data to parse cell-cell and gene-gene relationship governing mechanisms of cell identity.

## Supporting information

Supplemental Figures and tables description

## Data Availability

The mouse gastrulation atlas data is available in *MouseGastrulationData* [47] at https://github.com/MarioniLab/MouseGastrulationData. Code for TRIAGE-Cluster and TRIAGE-ParseR are available online at https://github.com/palpant-comp/TRIAGE-Cluster and https://github.com/palpant-comp/TRIAGE-ParseR. The combined analysis of the mouse gastrulation atlas data and *in vitro* iPSC differential data can be explored at http://cellfateexplorer.d24h.hk/.

## Author Contributions

Y.S. further developed, implemented, and benchmarked TRIAGE-Cluster, developed topographical visualisation approach, and wrote the manuscript. W.J.S. conceived and developed TRIAGE-ParseR and wrote the manuscript. S.S. helped develop TRIAGE-Cluster method. E.S. generated the overall pipeline Figure 1. D.P. and Z.S. constructed the website. D.M. performed pseudotime analysis of TRIAGE genes. M.D.W. interpreted the genetic basis of peak classifications in Figure 6. Q.N. and J.W.K.H. supervised website development. M.B. supervised TRIAGE-ParseR development and edited the manuscript. N.J.P. conceptualised the study, supervised the project, raised funding, and wrote the manuscript.

## Funding

Funding support was provided from the National Health and Medical Research Council of Australia (Grants 1143163 (N.P.), GNT2008928 (Q.N.), APP2013027 (M.W.)), the Australian Research Council (Grant SR1101002 (N.P.), DP220101878 (M.W.), FT200100899 (M.W.)), the National Heart Foundation of Australia (Grants 101889 and 106721, N.P.), the Medical Research Future Fund (APP2016033, N.P.), and AIR@InnoHK administered by Innovation and technology Commission (J.W.K.H.).

## Declarations of Interests

The authors declare no competing interests.

## Notes

### Competing Interest Statement

The authors have declared no competing interest.

### Summary of Updates

author name is updated; main text is updated; all figures are updated to match languages in main text.

