## Supplemental Figures and tables description for "Inferring cell diversity in single cell data using consortium-scale epigenetic data as a biological anchor for cell identity"

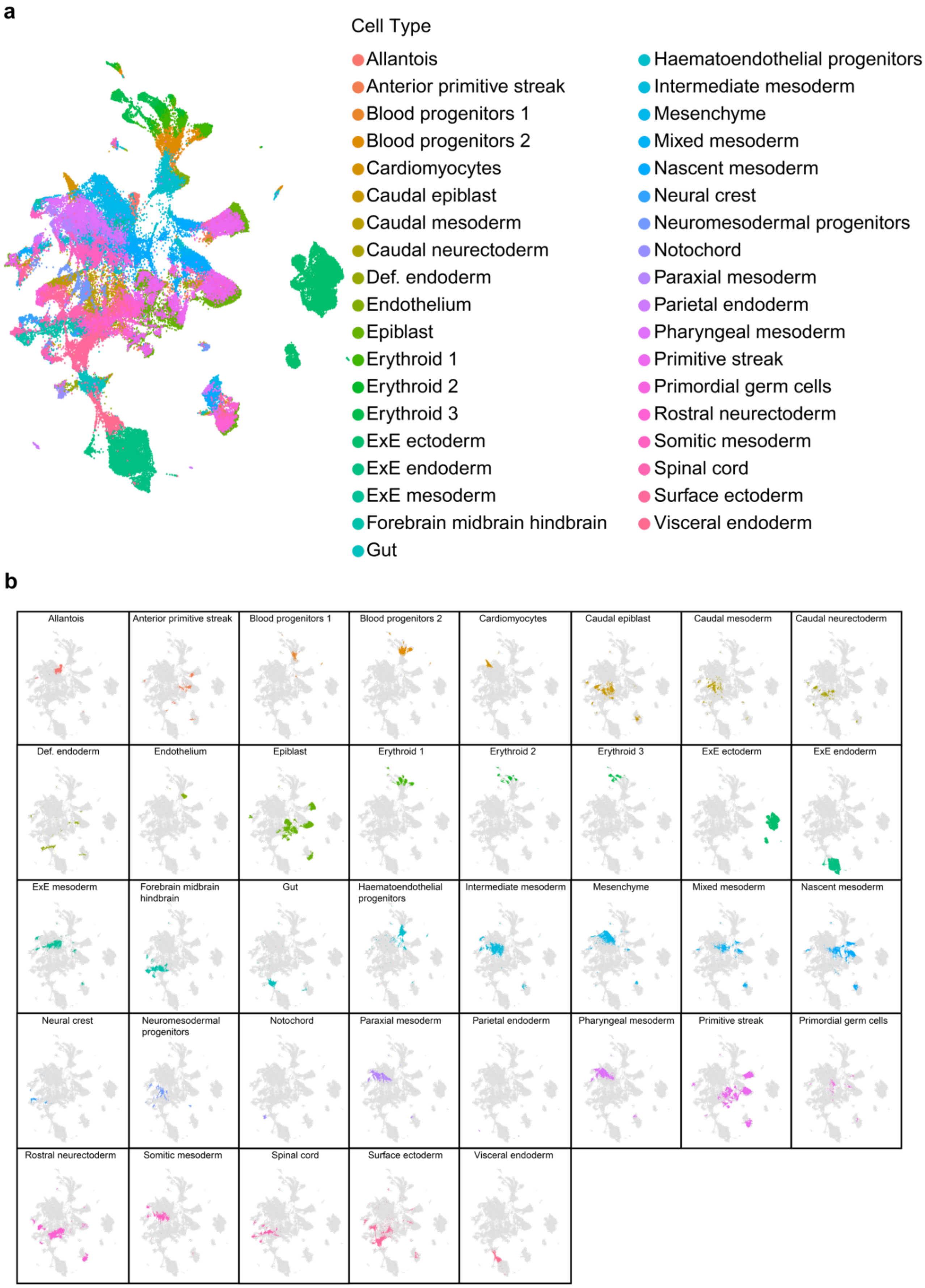


**Figure S1. Overview of original annotation for the mouse gastrulation atlas data.**

**a-b)** Original annotations are shown in single UMAP and separate in multiple UMAP plots.

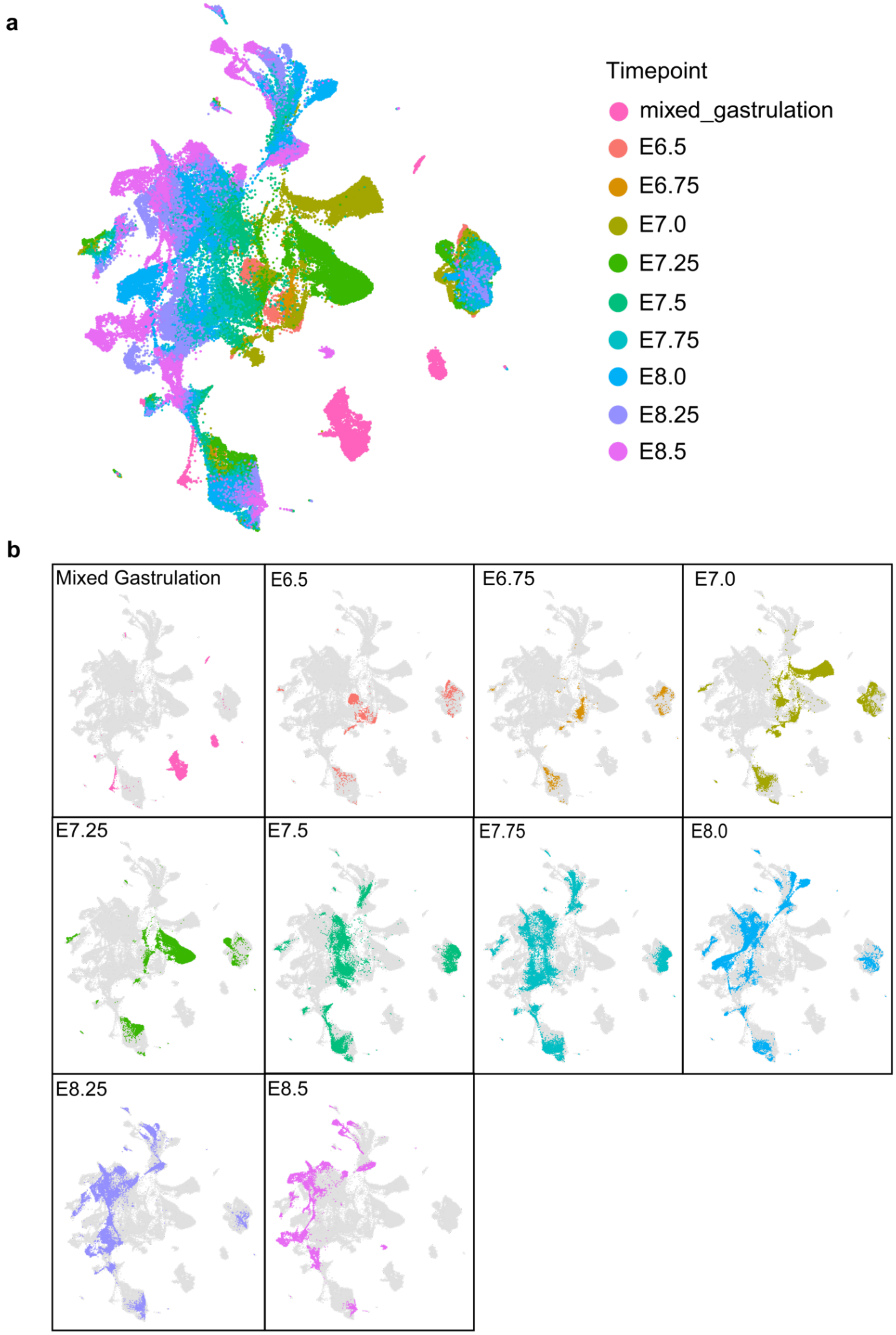


**Figure S2. Overview of time point for the mouse gastrulation atlas data.**

**a-b)** Time points are shown in single UMAP and separate in multiple UMAP plots.

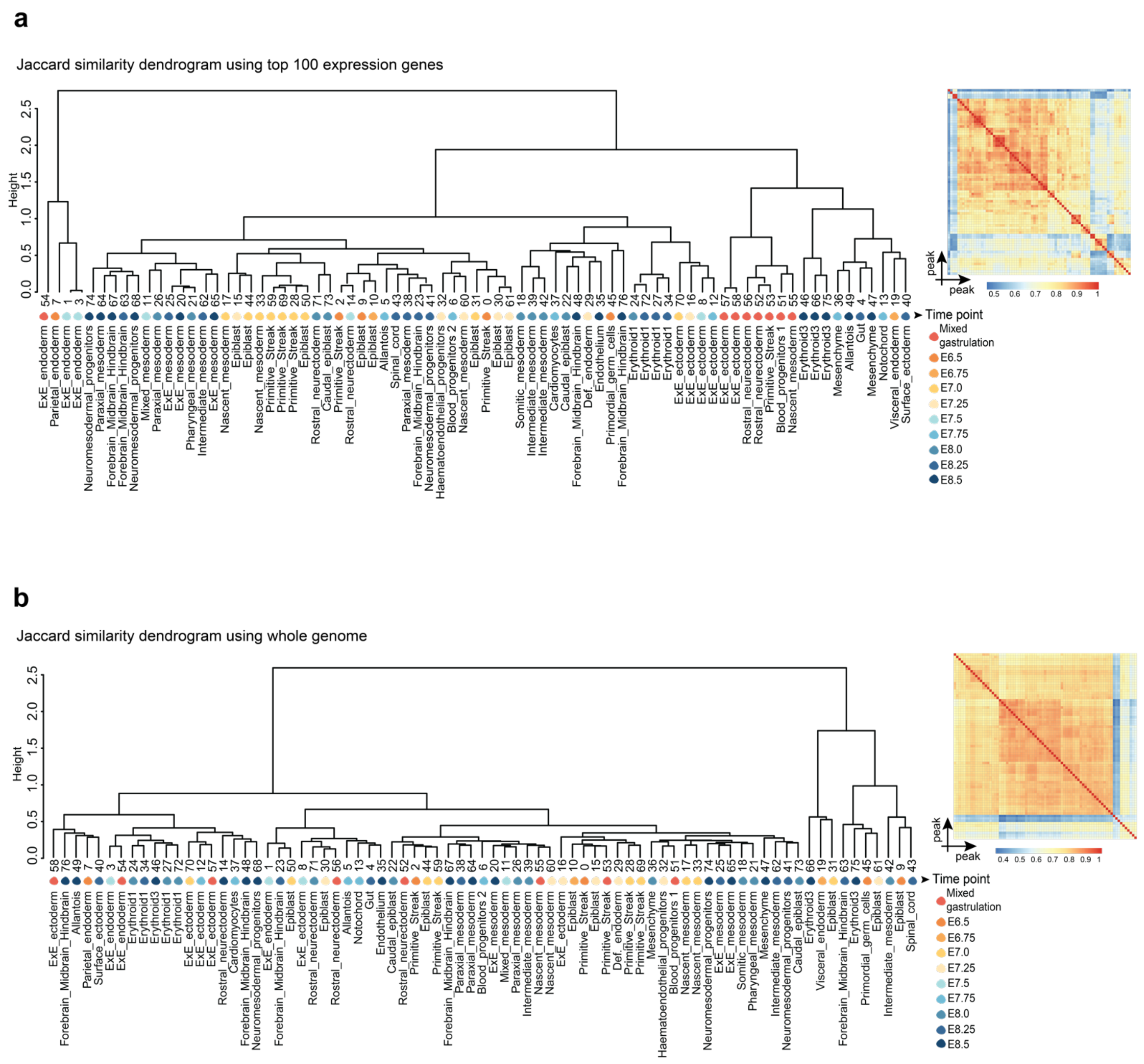


**Figure S3. Comparing cell type identification using different gene lists.**

**a-b)** Jaccard similarity dendrogram using top 100 genes ranked by original expression **(a)** and all genes expressed in each peak **(b)** at bandwidth 0.3 showing peak relationship (first row), mapped timepoint (second row), anchor gene (third row), and mapped original annotation (forth row).

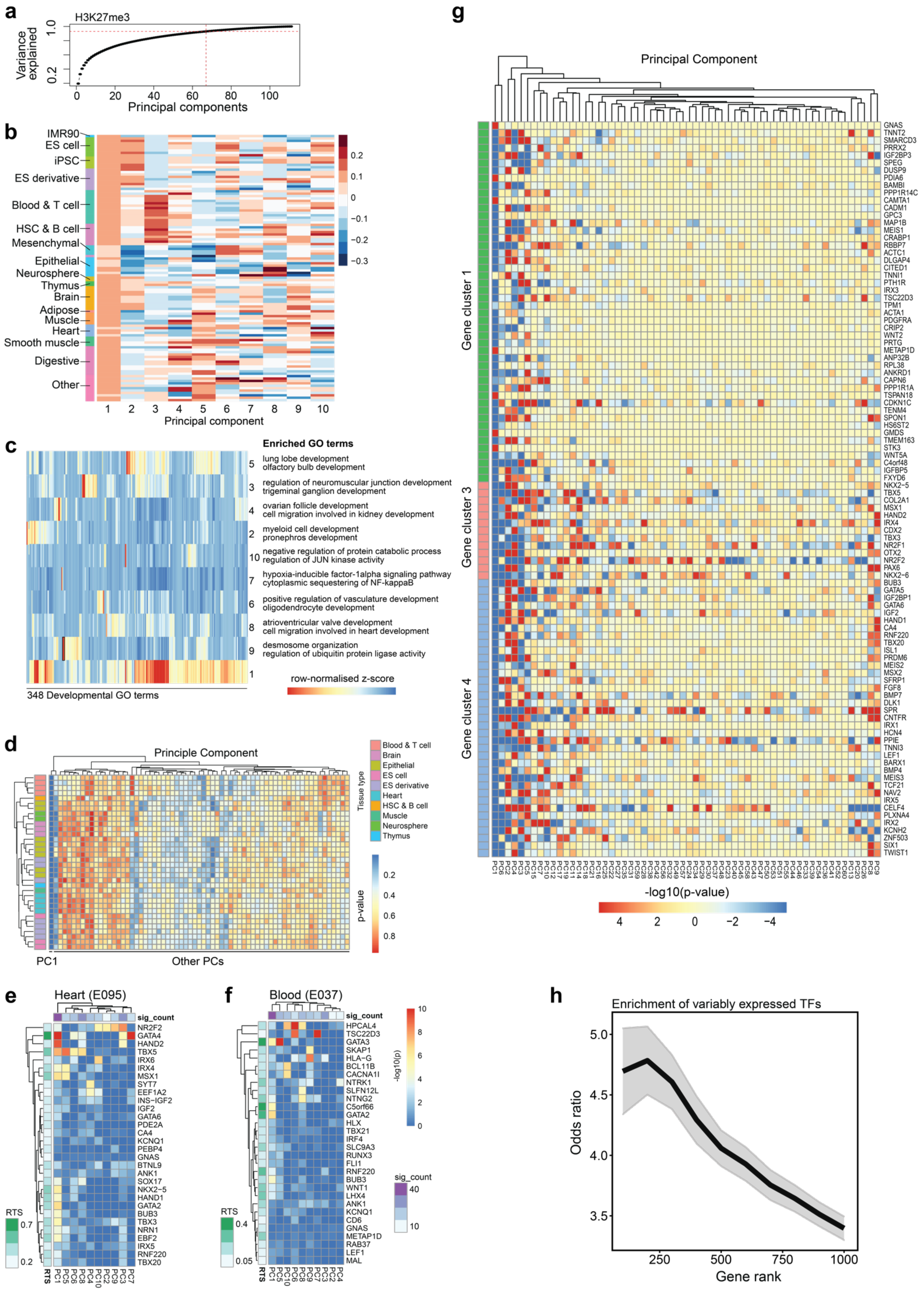
a
H3K27me3
Variance explained
Principal components
b
IMR90
ES cell
iPSC
ES derivative
Blood & T cell
HSC & B cell
Mesenchymal
Epithelial
Neurosphere
Thymus
Brain
Adipose
Muscle
Heart
Smooth muscle
Digestive
Other
Principal component
c
Enriched GO terms
lung lobe development
olfactory bulb development
regulation of neuromuscular junction development
trigeminal ganglion development
ovarian follicle development
cell migration involved in kidney development
myeloid cell development
pronephros development
negative regulation of protein catabolic process
regulation of JUN kinase activity
hypoxia-inducible factor-1alpha signaling pathway
cytoplasmic sequestering of NF-kappaB
positive regulation of vasculature development
oligodendrocyte development
atrioventricular valve development
cell migration involved in heart development
desmosome organization
regulation of ubiquitin protein ligase activity
row-normalised z-score
348 Developmental GO terms
d
Principle Component
Tissue type
Blood & T cell
Brain
Epithelial
ES cell
ES derivative
Heart
HSC & B cell
Muscle
Neurosphere
Thymus
p-value
PC1
Other PCs
e
Heart (E095)
sig_count
NR2F2
GATA4
HAND2
TBX5
IRX6
IRX4
MSX1
SYT7
EEF1A2
INS−IGF2
IGF2
GATA6
PDE2A
CA4
KCNQ1
PEBP4
GNAS
BTNL9
ANK1
SOX17
NKX2−5
HAND1
GATA2
BUB3
TBX3
NRN1
EBF2
IRX5
RNF220
TBX20
RTS
f
Blood (E037)
sig_count
HPCAL4
TSC22D3
GATA3
SKAP1
HLA−G
BCL11B
CACNA1I
NTRK1
SLFN12L
NTNG2
C5orf66
GATA2
HLX
TBX21
IRF4
SLC9A3
RUNX3
FLI1
RNF220
BUB3
WNT1
LHX4
ANK1
KCNQ1
CD6
GNAS
METAP1D
RAB37
LEF1
MAL
RTS
-log10(p)
sig_count
g
Principal Component
Gene cluster 1
Gene cluster 3
Gene cluster 4
GNAS
TNNT2
SMARCD3
PRRX2
IGF2BP3
SPEG
DUSP9
PDIA6
BAMBI
PPP1R14C
CAMTA1
CADM1
GPC3
MAP1B
MEIS1
CRABP1
RBBP7
ACTC1
DLGAP4
CITED1
TNNI1
PTH1R
IRX3
TSC22D3
TPM1
ACTA1
PDGFRA
CRIP2
WNT2
PRTG
METAP1D
ANP32B
RPL38
ANKRD1
CAPN6
PPP1R1A
TSPAN18
CDKN1C
TENM4
SPON1
HS6ST2
GMDS
TMEM163
STK3
WNT5A
C4orf48
IGFBP5
FXYD6
NKX2−5
TBX5
COL2A1
MSX1
HAND2
IRX4
CDX2
TBX3
NR2F1
OTX2
NR2F2
PAX6
NKX2−6
BUB3
GATA5
IGF2BP1
GATA6
IGF2
HAND1
CA4
RNF220
TBX20
ISL1
PRDM6
MEIS2
MSX2
SFRP1
FGF8
BMP7
DLK1
SPR
CNTFR
IRX1
HCN4
PPIE
TNNI3
LEF1
BARX1
BMP4
MEIS3
TCF21
NAV2
IRX5
CELF4
PLXNA4
IRX2
KCNH2
ZNF503
SIX1
TWIST1
-log10(p-value)
h
Enrichment of variably expressed TFs
Odds ratio
Gene rank

**Figure S4. H3K27me3 patterns mark genes regulating specific biological process.**

**a)** Principal component analysis (PCA) showing approximately 96.5% H3K27me3 variance can be explained by top 67 PCs in Roadmap data (red dashed line).

**b)** The heatmap shows enrichment of the genes with top 100 discordance score from each sample in Roadmap in each PC.

**c)** The heatmap shows the Gene Ontology (GO) functional enrichment of 348 terms in each PC.

**d)** Enrichment of H3K27me3 patterns among top 100 RTS priority genes in Roadmap samples. The heatmap shows an enrichment p-value of a given pattern (or PC) using Fisher's exact test (one-tailed).

**e-f)** Enrichment of H3K27me3 patterns among top 30 RTS priority genes in left ventricle heart sample (E095) and blood sample (E037). Significant count is the number of genes among top 100 RTS priority genes showing significant enrichment of a given pattern. The heatmap shows the enrichment p-value (as -log10(p)) of the gene.

**g)** The heatmap shows an enrichment of H3K27me3 patterns among top 100 RTS priority genes using Fisher's exact test (one-tailed) in the three gene clusters detected in cardiomyocytes from the mouse gastrulation atlas data.

**h)** Enrichment of variably expressed TFs in top genes ranked by discordance score. Average odds ratio of variably expressed (defined by expression coefficient of variation>1) TFs across 46 different cell and tissue types from NIH Roadmap data were shown, with +/- standard error of the mean.

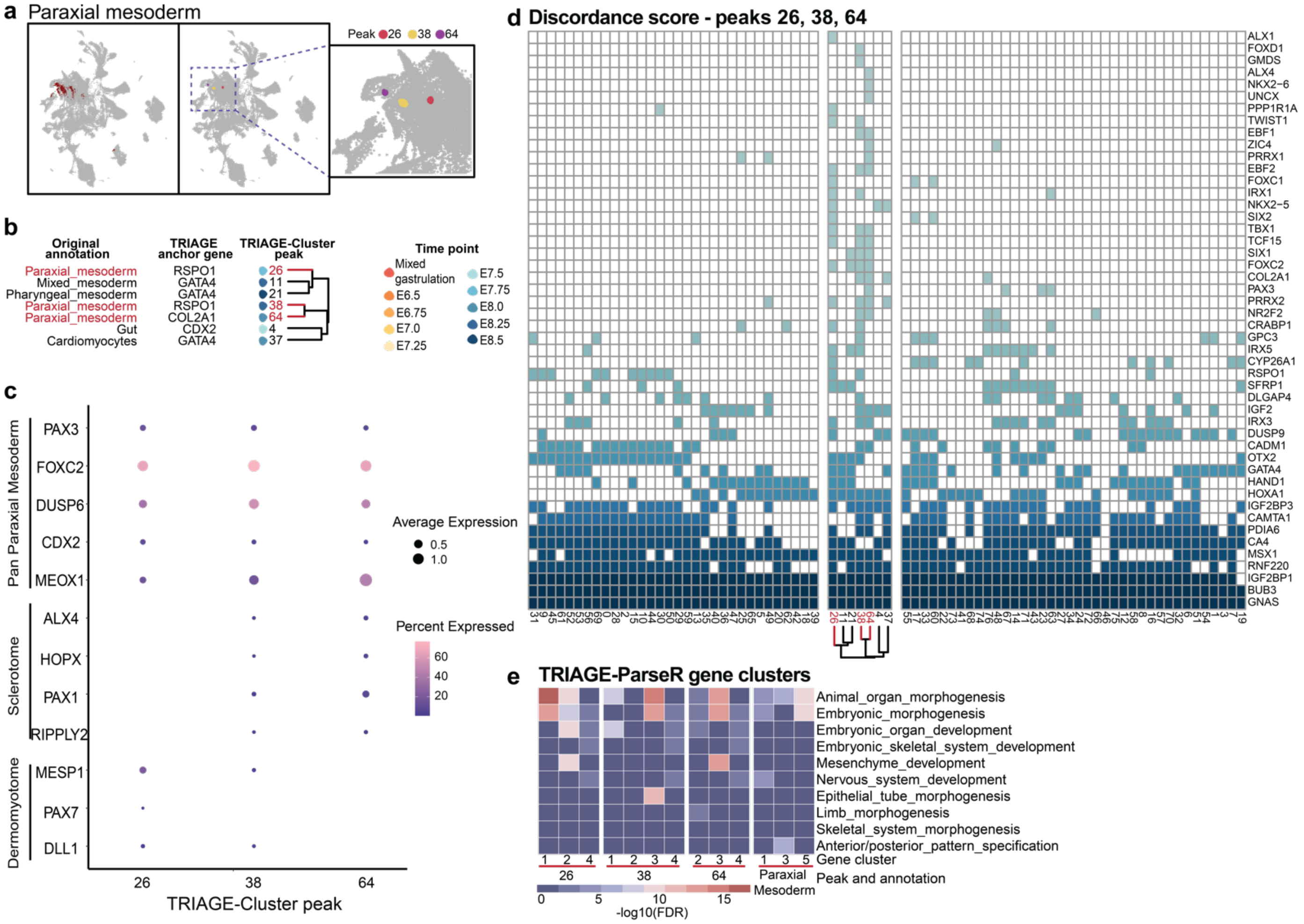


**Figure S5. TRIAGE-Cluster and TRIAGE-ParseR analysis enable identification of cell subtype diversity in single cell data.**

**a)** UMAP of original annotation paraxial mesoderm (left) and mapped TRIAGE-Cluster peaks (right and inset).

**b)** Jaccard similarity dendrogram (below) with bandwidth 0.3 showing three distinct "paraxial mesoderm" TRIAGE-Cluster peaks with their respective mapped timepoint and anchor genes (extracted from **Figure 2h**).

**c)** Dot plot showing marker genes associated with bone development lineage subtypes in each paraxial mesoderm peak.

**d)** Genes ranked by discordance score for each paraxial mesoderm peak (red) and measured across all other peaks. Output focuses on genes enriched in paraxial mesoderm peaks (row) for each TRIAGE-Cluster peak (column) to facilitate cell subtype classification.

**e)** Heatmap showing skeletal cell subtype gene ontology enrichment (-log10 (FDR)) from PCA-GMM analysis across gene clusters in the three paraxial mesoderm peaks (the first three blocks) and original paraxial mesoderm cells (the last block).

### Supplemental Tables

**Table S1. Gene RTS scores.**

**Table S2. Software details.**

**Table S3. EpiMap data.**

**Table S4. Gene Ontology for 17 peaks from TRIAGE-Cluster.**

**Table S5. Gene Ontology for 77 peaks from TRIAGE-Cluster.**

**Table S6. Gene Ontology for 17 peaks from Seurat clustering.**

**Table S7. Gene Ontology for 77 peaks from Seurat clustering.**
